# Immigration and establishment of urban *Trypanosoma cruzi* populations

**DOI:** 10.1101/515049

**Authors:** Alexander S.F. Berry, Renzo Salazar-Sánchez, Ricardo Castillo-Neyra, Katty Borrini-Mayorí, Claudia Chipana-Ramos, Melina Vargas-Maquera, Jenny Ancca-Juarez, César Náquira-Velarde, Michael Z. Levy, Dustin Brisson, Chagas Disease Working Group in Arequipa

## Abstract

Changing environmental conditions, including those caused by human activities, reshape biological communities through both loss of native species and establishment of non-native species in the altered habitats. Dynamic interactions with the abiotic environment impact both immigration and initial establishment of non-native species into these altered habitats. The repeated emergence of disease systems in urban areas worldwide highlights the importance of understanding how dynamic migratory processes affect the current and future distribution and abundance of pathogens in urban environments. In this study, we examine the pattern of invasion of *Trypanosoma cruzi*—the causative agent of human Chagas disease—in the city of Arequipa, Peru. Phylogenetic analyses of 136 *T. cruzi* isolates from Arequipa and other South American locations suggest that only one *T. cruzi* immigrant established a population in Arequipa as all *T. cruzi* isolated from vectors in Arequipa form a recent monophyletic group within the broader South American phylogeny. We discuss several hypotheses that may explain the limited number of established *T. cruzi* lineages despite multiple introductions of the parasite.

**Author Summary:** Human-associated pests and pathogens, who benefit from the abundance of humans and human-associated hosts or vectors, commonly invade environments altered by human activities. As the number and size of human-disturbed environments increase, so does the importance of identifying ecological and environmental factors that affect the probability that disease systems immigrate to, subsequently establish populations in, urban environments. We examined the number and timing of immigration and establishment events of *Trypanosoma cruzi*, the causative agent of Chagas disease, in a currently endemic area. Phylogenetic analyses of 136 *T. cruzi* isolates suggests that the current population descended from a single, recent immigration event. We discuss historical and ecological hypotheses that can explain the limited *T. cruzi* diversity in this region.

## Introduction

Habitat alterations are transforming biological communities worldwide [1–3]. The current and future geographic distributions of many species in disturbed environments depends upon their interactions with novel biotic and abiotic features during immigration and while establishing a growing population [4]. Although many species fail to establish thriving populations in altered habitats, others are well-suited to migrate to, and prosper in, these novel environments. For example, several populations of plant [5–7], insect [8–11], mammal [12,13], and bird [14,15] species are severely diminished or suffer local extinctions in recently urbanized environments [1], while several microbial species benefit from the abundance of humans and human-associated hosts or vectors in similar habitats [16]. Although conservation efforts have focused primarily on the impacts of environmental changes on native plant and animal species, establishment or population growth of disease-causing microbial populations can have a strong negative impact on populations of native flora and fauna [17] in addition to their impact on human health and economy [18]. The rate or impacts of invasions of infectious microbes may be mitigated through public health programs based on an understanding of the dynamic processes determining immigration and establishment rates. The regularity at which disease systems are emerging in many urban and urbanizing areas underscores the importance of understanding how disease-causing microbes migrate to, and establish in, urban environments [16], one of the most dramatic examples of habitat alteration [19,20]. In this study, we examine the patterns of invasion of *Trypanosoma cruzi*—the causative agent of Chagas disease in humans—into the city of Arequipa, Peru.

Invasion of a new environment by a pathogen occurs in three stages: (1) immigration, or the movement of an individual to the new environment; (2) establishment of a population via reproduction and population growth; and (3) local dispersal [4]. The many studies focusing on outbreaks of disease systems have generated a wealth of knowledge concerning factors affecting population growth [21,22] and considerable progress in understanding local dispersal [23]. For example, prior studies concluded that human-created containers increase the abundance of standing water that provide breeding habitats for the mosquitos that spread dengue virus [24]. Relatively few studies, by contrast, have investigated the early stages of invasion, the immigration and establishment processes, due to the practical difficulties of collecting the necessary data before a novel species is established.

The Chagas disease system in Arequipa, Peru, provides an ideal system in which to study the early invasion processes in urban environments. Arequipa has experienced rapid urbanization and human population growth in the previous half century with a concurrent population of *T. cruzi* [25]. The expansion in population sizes and geographic range of humans and their domestic animals provide hosts for both *T. cruzi* and its primary insect vector, *Triatoma infestans* [26–31]. The population history of the *T. cruzi* currently in Arequipa - including the geographic locations of the migrants that established the current population, the rate at which migrants enter and establish in the area, and the age of each established lineage - have not been investigated. Here, we performed phylogenetic analyses of maxicircle DNA, a non-recombining circular element analogous to mitochondrial DNA, to estimate the number of independent *T. cruzi* lineages established in Arequipa and to estimate the timing of each establishment event. We assessed whether extant *T. cruzi* in Arequipa form a single monophyletic clade, indicative of the establishment of a single migrant lineage, or multiple diverse clades, indicative of multiple independent immigration and establishment events.

## Results

The maxicircle sequence is useful for population genetic and phylogenetic analyses because it is conserved among diverse *T. cruzi* lineages and is likely non-recombining [32]. While there was substantial maxicircle sequence diversity among samples across South America, almost no diversity was observed among the samples from Arequipa. Over 13% (2055/15367bp) of the maxicircle sites were polymorphic among all 136 samples while only 16 sites were polymorphic (0.1%) among the 123 samples collected within Arequipa (Table 1). Similarly, estimates of diversity among the samples from Arequipa derived from population genetic statistics were substantially lower than the total diversity across all samples (π=6.8*10^-5^ vs 8.18*10^-3^; θ=1.93*10^-4^ vs 2.44*10^-2^; average pairwise distance 1.04 vs 126). In contrast to the limited genetic diversity within Arequipa, other locales from which multiple isolates were sampled contain substantial genetic diversity among limited samples (N<3; Fig 1).

**Table 1.** Population Genetic Statistics.

| | Average Pairwise Distance | $\pi$ | $\theta$ | Segregating Sites |
| --- | --- | --- | --- | --- |
| All samples (N=136) | 126 | $8.18*10^{-3}$ | $2.44*10^{-2}$ | 2055 |
| Arequipa (N=123) | 1.04 | $6.80*10^{-5}$ | $1.93*10^{-4}$ | 16 |
| South America (N=13) | 728 | $4.73*10^{-2}$ | $4.24*10^{-2}$ | 2022 |
| La Esperanza (N=3) | 643 | $4.18*10^{-2}$ | $4.18*10^{-2}$ | 964 |

**Figure 1.**
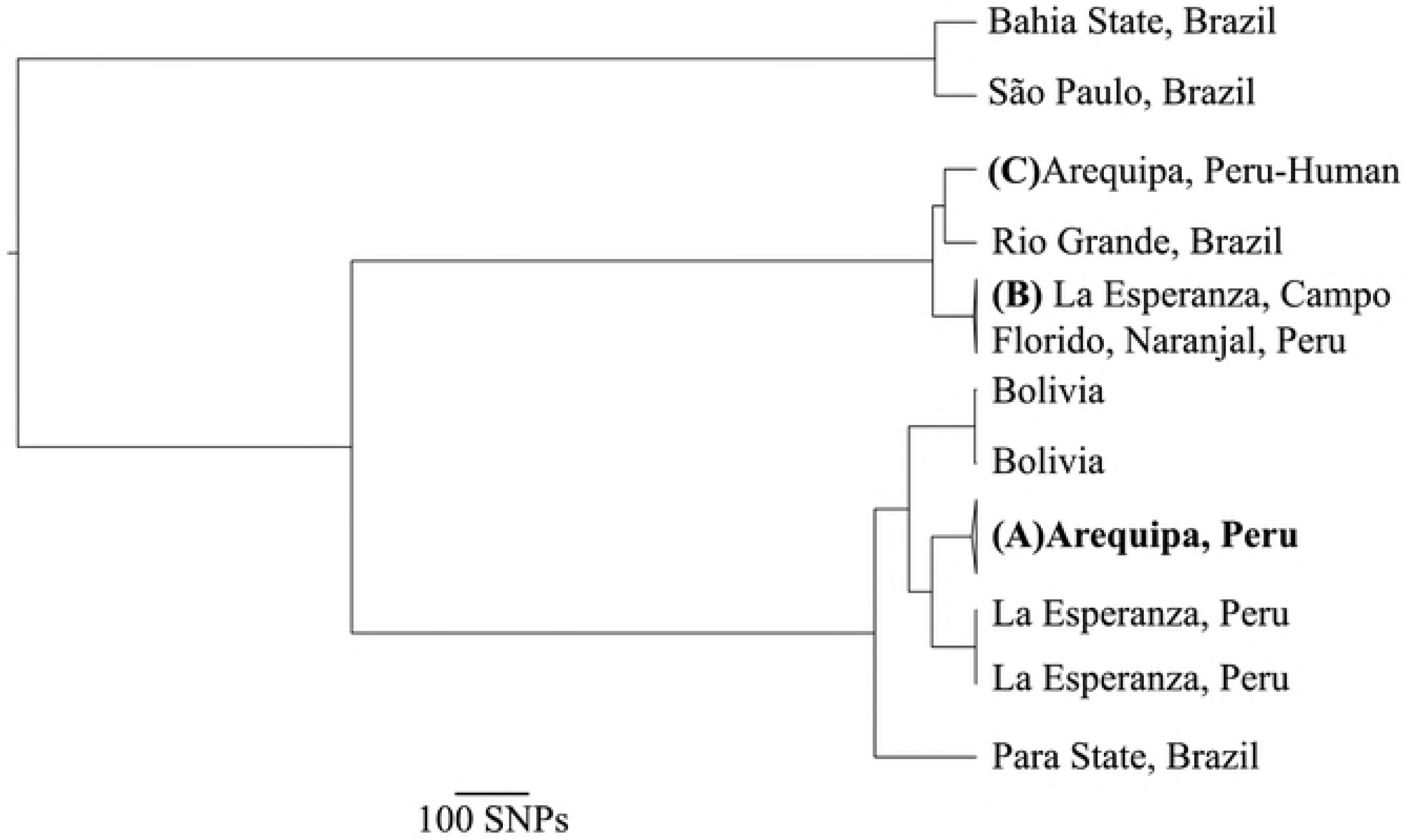
The extant *T. cruzi* population in Arequipa arose from a single, recent introduction. Maximum clade credibility (MCC) tree shows that **(A)** all 123 *T. cruzi* isolated from bugs and domestic mammals in Arequipa form a monophyletic clade with a single, recent common ancestor, indicative of a single immigration event in the recent past. Despite substantial genetic diversity among *T. cruzi* throughout South America, those collected in Arequipa show little diversity. **(B)** Three samples collected in Campo Florido and Naranjal, Peru and one sample from La Esperanza, Peru have nearly identical maxicircle sequences and form a monophyletic clade. La Esperanza, Peru contains at least two distinct *T. cruzi* lineages, suggesting multiple independent introductions. **(C)** The only *T. cruzi* sample isolated from a human in Arequipa is distinct from all other samples from Arequipa, suggesting that this introduction has not established in the city. All tips represent a single sample except (A) (N=123) and (B) (N=4). All nodes have strong support (posterior probability≥0.99). Nodes are collapsed when the samples contained have nearly identical maxicircle sequences.

The monophyletic group containing all 123 samples derived from *T. infestans* and domestic animals collected in Arequipa coalesce in the very recent past, despite collection sites extending throughout Arequipa and surrounding towns across 7 years (Fig 1). One sample derived from an infected human in Arequipa (Fig 1C) belongs to a lineage that is more closely related to samples collected in Rio Grande, Brazil than to the other samples collected in Arequipa (Fig 1). This sample is distinct from all other samples in Arequipa suggesting that *T. cruzi* can immigrate to Arequipa but may not establish in the vector population. The population size of an unsampled *T. cruzi* lineage – if present in the *T. infestans* population – must be at least 42 times smaller than the dominant population in Arequipa to have remained undetected by chance (p<0.05).

In contrast to the monophyletic ancestry found in Arequipa, genetic diversity was apparent in the samples collected from other regions, despite limited numbers of samples (N<3). Population genetic diversity among the three samples collected La Esperanza, a town of 57 houses in the Cutervo Province of Cajamarca, Peru (π=4.18*10^-2^; θ=4.18*10^-2^), are much larger than those in Arequipa despite the limited number of samples [33]. There is no statistical correlation between genetic relatedness and geographic distance among South American samples (Table 2). For example, while proximal towns La Esperanza and Campo Florido, Peru have closely related *T. cruzi*, isolates from cities around Brazil encompass nearly the total genetic diversity.

**Table 2.** Distance matrix showing average pairwise SNP distance between samples (bottom triangle) and Euclidean distance between sample collection locations (top triangle).

|  | (A)<br>Arequipa,<br>Peru | La<br>Esperanza,<br>Peru | La<br>Esperanza,<br>Peru | (B) La<br>Esperanza,<br>Campo Florido,<br>& Naranjal, Peru | Bolivia | Bolivia | (C) Arequipa,<br>Peru - Human | São Paulo,<br>Brazil | Para State,<br>Brazil | Bahia State,<br>Brazil | Rio Grande,<br>Brazil |
| --- | --- | --- | --- | --- | --- | --- | --- | --- | --- | --- | --- |
| (A)<br>Arequipa,<br>Peru |  | 1400km | 1400km | 1400km | 700km | 700km | 0km | 2700km | 2500km | 3200km | 2500km |
| La<br>Esperanza,<br>Peru | 95 |  | 0km | 60km | 2000km | 2000km | 1400km | 4000km | 3000km | 4100km | 3900km |
| La<br>Esperanza,<br>Peru | 95 | 0 |  | 60km | 2000km | 2000km | 1400km | 4000km | 3000km | 4100km | 3900km |
| (B) La<br>Esperanza,<br>Campo Florido,<br>& Naranjal, Peru | 970 | 966 | 966 |  | 2000km | 2000km | 1400km | 4000km | 3000km | 4100km | 3900km |
| Bolivia | 137 | 137 | 137 | 957 |  | 0km | 700km | 2000km | 1800km | 2500km | 2000km |
| Bolivia | 137 | 137 | 137 | 958 | 2 |  | 700km | 2000km | 1800km | 2500km | 2000km |
| (C) Arequipa,<br>Peru - Human | 975 | 971 | 971 | 89 | 967 | 968 |  | 2700km | 2500km | 3200km | 2500km |
| São Paulo,<br>Brazil | 1044 | 1045 | 1045 | 1033 | 1045 | 1046 | 1058 |  | 2100km | 1300km | 1100km |
| Para State,<br>Brazil | 216 | 216 | 216 | 984 | 205 | 205 | 992 | 1059 |  | 1500km | 2800km |
| Bahia State,<br>Brazil | 1284 | 1289 | 1289 | 1234 | 1287 | 1287 | 1258 | 68 | 1308 |  | 2500km |
| Rio Grande,<br>Brazil | 1005 | 1003 | 1003 | 82 | 1001 | 1001 | 77 | 1052 | 1026 | 1284 |  |

(A) 123 samples isolated from bugs, dogs, and guinea pigs in Arequipa are represented here.
(B) 4 samples collected in Campo Florido, Naranjal, and La Esperanza are represented here.
Euclidean distances (top triangle) are displayed for Santa Cruz.
(C) The only sample isolated from a human in Arequipa is represented here.

## Discussion

The invasion of recently altered environments by non-native species impacts the population health of native species as well as human health and economy. Investigation of the dynamic process of immigration and establishment of non-native species into these disturbed habitats has the potential to mitigate the impacts of pests and pathogens that are detrimental to human populations as well as agricultural and native species [17]. The analyses presented here suggest that the population of *T. cruzi* in Arequipa, Peru, descended from a single recent invasion. This conclusion is supported by the extremely limited genetic diversity observed among *T. cruzi* isolates sampled within and around the city, in contrast with considerable genetic diversity observed regionally and at other locales (Table 1; Fig 1). Several non-exclusive hypotheses may explain these results, including a low immigration rate, that immigration is common but immigrants rarely establish populations as a result of the low transmission rate between hosts and vectors, or that there is a high turnover rate among *T. cruzi* lineages.

Successful invasion of a novel geographic area is a function of the rate of immigration, the temporal duration that a habitat has been suitable for establishment, and the probability that an immigrant can reproduce and establish a population. The influx of humans and associated products and domesticated animals into Arequipa over the last ~60 years due to rapid urbanization and economic growth [25,34] has provided many opportunities for *T. cruzi* immigration. However, many migrants to Arequipa come from areas in which they were not exposed to *T. cruzi*, such as the neighboring regions of Puno and parts of Cusco which are beyond the range of *T. infestans* [35]. Nevertheless, one *T. cruzi* lineage that does not appear to be circulating within the *T. infestans* population was detected in an infected human (Fig. 1), suggesting that *T. cruzi* immigration through human movement can occur. These data suggest that multiple *T. cruzi* lineages may have immigrated to Arequipa with all but one failing to transmit sufficiently to establish a population. While the data presented here are consistent with the hypothesis that *T. cruzi* in Arequipa originated in Bolivia [36,37], future studies will be necessary to identify the source, rate, and potential mechanism of *T. cruzi* immigration, independent of establishment probability, through analyses of the genomic diversity in human infections.

Prior studies across multiple species suggest that the majority of immigrants in most species that reach a novel geographic area fail to establish a population due to both inhospitable local environmental conditions [1,38] and stochasticity [39,40]. Environmental factors that can reduce establishment probabilities include unfavorable abiotic conditions, limited food resources or vectors, or an abundance of predators or competitors. The establishment probability of immigrant *T. cruzi* in an urban environment is likely depressed by a low transmission rate from infected humans to vectors [41–44]. For example, many human immigrants moved to locations in the city without established *T. infestans* populations [25,35], which may have resulted in few opportunities for *T. cruzi* transmission from infected immigrant humans to insect vectors, thus curtailing establishment probabilities. Further, the transmission probability of *T. cruzi* from *T. infestans* that do acquire the parasite to novel hosts is low [42,45,46], thus reducing the probability that a recently-immigrated *T. cruzi* lineage will establish a population. Both a low probability of establishing a population due to limited contact between *T. infestans* and infected immigrant humans or due to the limited probability of transmission from vectors to host are consistent with the observation of only a single established *T. cruzi* lineage in vectors in Arequipa.

The observed establishment probability is likely independent of competition among *T. cruzi* lineages. Competitive exclusion—where an existing population prevents the invasion of new immigrants—appears unlikely as the majority of city blocks do not contain *T. cruzi* [25,47] despite substantial vector populations [48]. Under the competitive exclusion hypothesis, one might expect different *T. cruzi* lineages establishing in different areas of the city. Indeed, multiple *T. cruzi* lineages do co-circulate within the same locality [44,49–53], as seen in the samples sequenced here from La Esperanza (Fig 1), and even within the same host [54,55], suggesting that competition is not preventing the establishment of multiple lineages in Arequipa.

The limited genetic diversity within Arequipa could result from a recent replacement of a previously dominant lineage through natural population processes. While the continuous substitution of a dominant strain through natural selection or drift is common in well-mixed populations, geographic structure within populations tends to result in the persistence of genetic diversity [56]. The absence of samples deriving from a previously established *T. cruzi* lineage in the fragmented urban and inter-district landscapes, much of which contains an active vector population but no *T. cruzi* [25,47,48], is suggestive that no previous lineages dominated this area. Additionally, no other *T. cruzi* lineage circulating in the vector population was detected despite the temporal range of samples in our dataset (7 years).

In conclusion, all relevant data suggest that the vast majority of, if not all, *T. cruzi* circulating in vector populations prior to the recently-controlled epidemic in the city of Arequipa descended from a single immigrant. The single divergent lineage found in a human patient suggests that *T. cruzi* may regularly immigrate to the city but that immigrants rarely establish populations. Low genetic diversity, coupled with minimal gene flow into the city, could limit the phenotypic diversity that results in variable effectiveness of treatment options and diagnoses regularly associated with Chagas disease [57,58]. However, a potential downside of the limited diversity is that many currently available diagnostics may be ineffective against the local lineage [59,60], as seen in Arequipa [61]. It may be necessary to use locally-optimized diagnostics in endemic regions, which highlights the difficulties in diagnosing *T. cruzi* infection in immigrants to non-endemic countries such as the United States [62,63].

## Methods

### Ethics Statement

The Institutional Animal Care and Use Committee (IACUC) of Universidad Peruana Cayetano Heredia reviewed and approved the animal-handling protocol used for this study (identification number 59605). The IACUC of Universidad Peruana Cayetano Heredia is registered in the National Institutes of Health at the United States of America with PHS Approved Animal Welfare Assurance Number A5146-01 and adheres to the Animal Welfare Act of 1990 [64].

### Sample collection

DNA from 133 *T. cruzi* isolates collected over 7 years (2008 to 2015) were analyzed to determine phylogenetic relationships (Fig 2; Fig 3; Fig S1). The majority of samples were isolated from *T. infestans* bugs collected from houses throughout Arequipa (N=114). Three of these samples were obtained using xenodiagnosis as described in Chiari & Galvão (1997). An additional ten samples from Arequipa were isolated from the blood of guinea pigs (N=7), dogs (N=2), and a human (N=1). Six isolates were derived from *Panstrongylus lignarius* (N=5) - also known as *P. herreri* [65] - and one guinea pig (N=1) collected in the small towns of La Esperanza, Campo Florido, and Naranjal in northern Peru [33]. Cultures of three previously established strains isolated from humans in Bolivia (Bol-SH001 and Bol-DH29) and São Paulo, Brazil (TC-y) were provided by the Infectious Diseases Research Laboratory at Universidad Peruana Cayetano Heredia (Fig 3).

**Figure 2.**
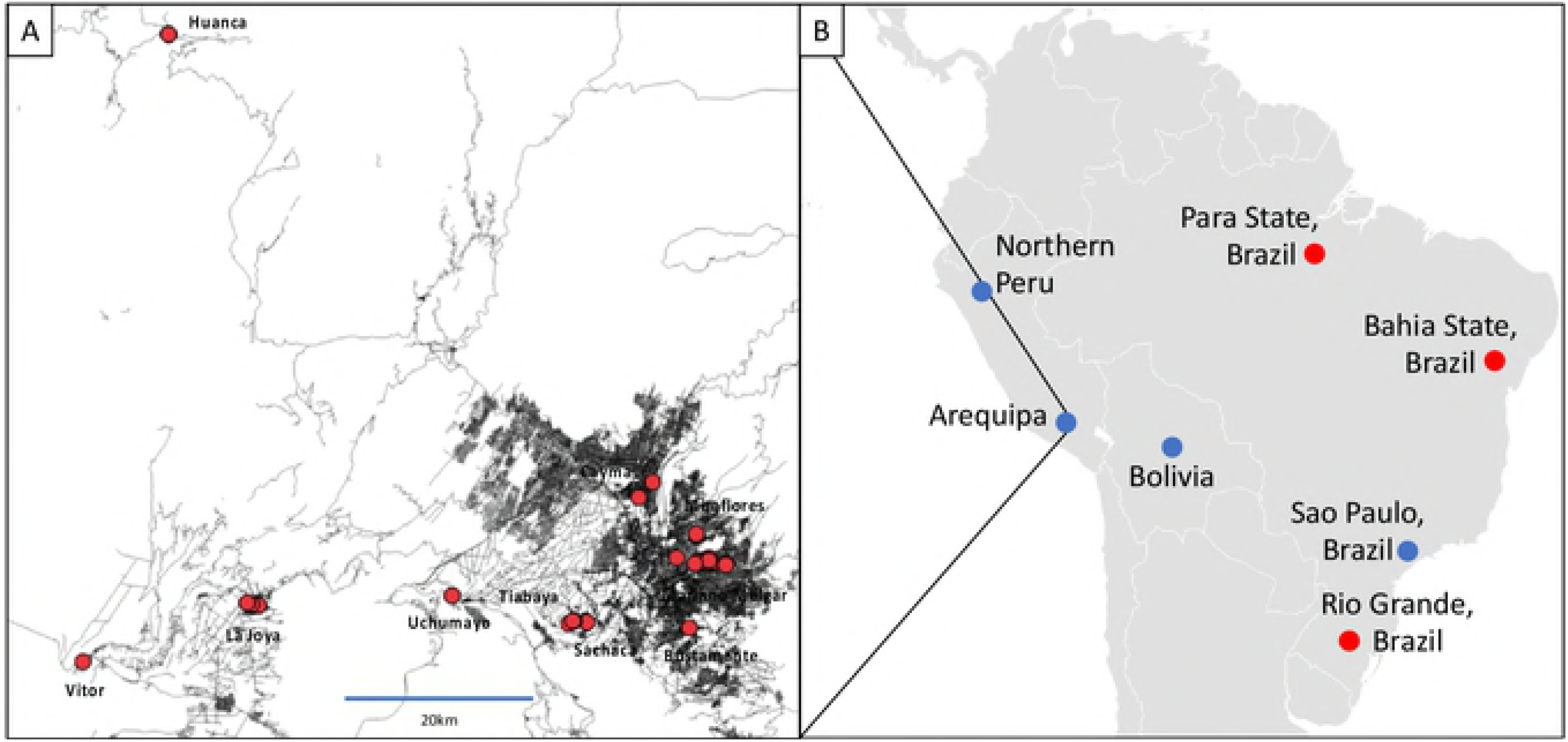
Spatial distribution of samples collected in (A) Arequipa, Peru and (B) South America. **(A)** The names of ten districts from which *T. cruzi* samples were collected are labeled. Houses from which isolates were collected are represented by red dots. Lines represent major roadways. Densely populated areas appear grey due to the density of roads. Map of Arequipa with sample locations was generated using QGIS v. 2.1 [80]. **(B)** The sites where isolates were collected are represented by blue dots. Neighboring towns of La Esperanza, Campo Florido, and Naranjal are represented by a single blue dot labeled “Northern Peru”. Sequences obtained from NCBI database are represented by red dots. Map of South America was modified from https://commons.wikimedia.org/wiki/Atlas_of_the_world.

**Figure 3.**
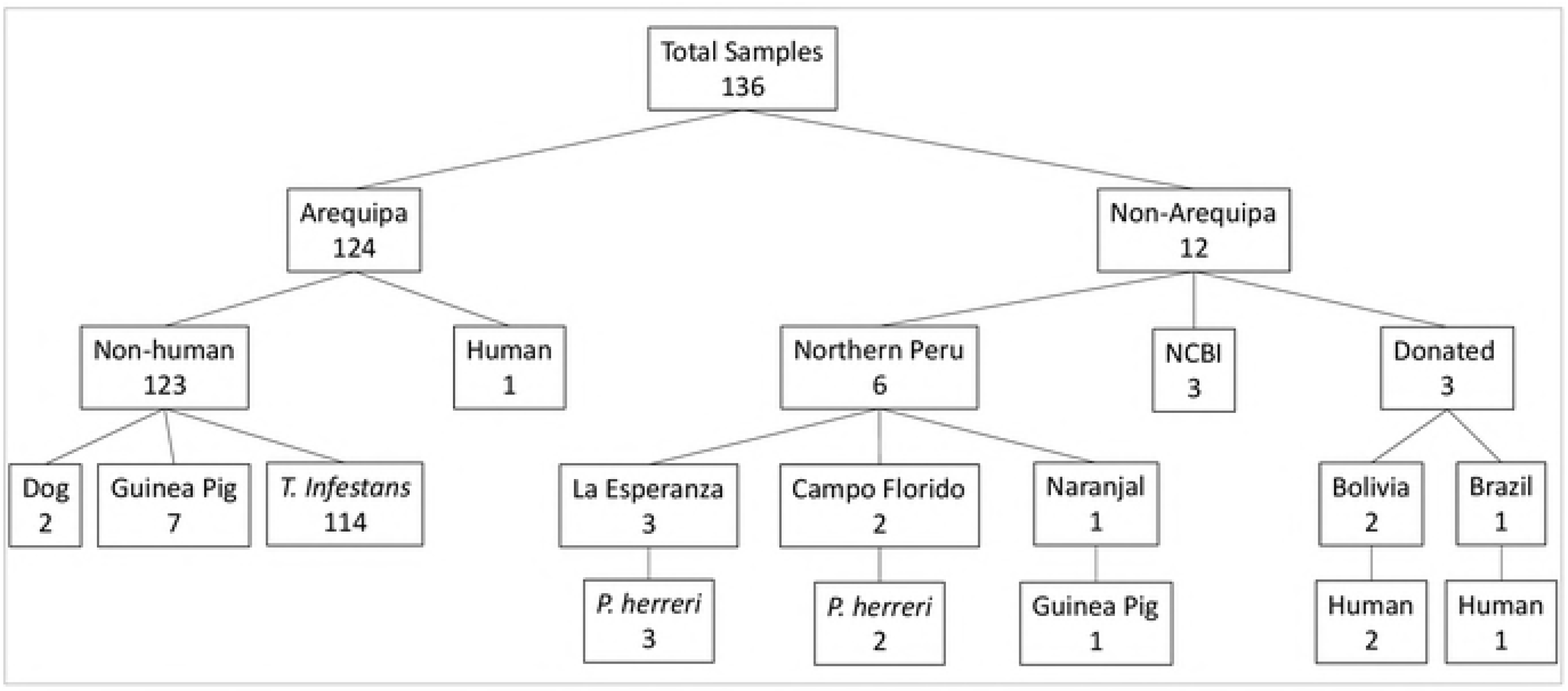
Number of samples collected from each host species per location. Most samples (N=124) were collected in Arequipa and 6 from small towns in northern Peru. 3 isolates were provided by the Infectious Diseases Research Laboratory at Universidad Peruana Cayetano Heredia. Maxicircle sequences for 3 *T. cruzi* lineages were downloaded from the NCBI database.

*T. cruzi* was isolated from vertebrates (N=8) using an adaptation of an artificial feeding system that was originally described in Harington (1960) [66]. Briefly, each blood sample was collected with citrate-phosphate-dextrose, transferred into a small plastic jar, and covered with a latex membrane fitted tightly with a rubber band. The jars were placed into an incubator and gradually heated to 35° C. Once the temperature was reached, the jars were inverted to allow uninfected *T. infestans* to feed through the membrane for 15 minutes. *T. cruzi* from the eight laboratory-infected *T. infestans*, all 114 naturally-infected *T. infestans*, and five naturally-infected *P. lignarius* were passaged through guinea pigs or mice in order to avoid isolating other microbes present in the vector, as described in Castillo-Neyra *et al.* (2016) [67]. Feces from infected vectors were injected into guinea pigs or mice and *T. cruzi* was isolated from the blood of each experimentally-infected mammal. *T. cruzi* were directly isolated in LIT culture media from the blood samples of three naturally-infected guinea pigs collected in Arequipa without passage through *T. infestans.*

Reference sequences of three *T. cruzi* isolates obtained from NCBI database were used in subsequent analyses: Silvio, isolated from a human in Para State, Brazil; Esmeraldo, isolated from a human in Bahia State, Brazil; and CL Brener, isolated from a human in Rio Grande, Brazil [68] (Fig 3).

### Sequencing

DNA from all laboratory cultures was extracted using Qiagen DNEasy DNA Purification Kit. 150bp single-end read libraries were prepared using TruSeq Nano kit and sequenced to an average depth of >50X using Illumina’s NextSeq500. Six *T. cruzi* isolates were prepared in duplicate, and one in triplicate, to allow estimation of sequencing error. Low quality bases were trimmed from raw reads using trimmomatic-0.32 [69].

### Sequence assembly

Bowtie2 [70] was used to assemble maxicircle sequences to the most closely related reference sequence, Silvio (gi|225217165|gb|FJ203996.1), obtained from NCBI [68]. Duplicate reads were removed from the assembly using Picard’s MarkDuplicates functionality [71]. The assembly had an average depth of >600X across all maxicircles. Maxicircle consensus sequences were determined using VarScan [72], ensuring highly-confident base calls by requiring a 60% match to call each SNP.

### Maxicircle alignment

All assembled maxicircle sequences and the reference were aligned to the Silvio partial maxicircle sequence (gi|225217165|gb|FJ203996.1|), Esmeraldo strain complete maxicircle (gi|85718082|gb|DQ343646.1), and the CL Brener complete maxicircle (gi|85718081|gb|DQ343645.1) downloaded from the NCBI database. The sequences were aligned using MUSCLE as implemented in MEGA7 [73]. The ends were trimmed so that all sequences started and ended on the same nucleotide, resulting in a final alignment of 15357bp.

### Phylogenetic analyses

Phylogenetic analyses of the 15357bp maxicircle sequence from all samples and reference strains were performed using BEAST 1.8.4 [74]. Phylogenetic analyses assumed a model of sequence evolution in which the rates of A→T, C→G, and G→T are equal (123343) with γ-distributed rate heterogeneity. An Extended Bayesian Skyline tree prior [75] with constant evolutionary rates across lineages (strict clock) was chosen based on BEAST Model Test implemented in BEAST2 [76]. Starting with a UPGMA tree and running one Markov chain Monte-Carlo chain for each of five independent runs of 20 million iterations sampling every 2000 iterations ensured sufficient mixing after a 10% burn-in (ESS values >200 in Tracer v1.6.0) [77]. Tree files were combined using LogCombiner1.8.4, excluding a 10% burn-in for each. A Maximum Clade Credibility tree was generated from the combined tree file using TreeAnnotator 1. 8.4 and FigTree v1.4.2 was used to visualize tree files (available at http://beast.bio.ed.ac.uk). Phylogenetic analyses were performed using the BEAGLE library to increase computation speed [78,79].

### Statistical Analyses

Metrics of population genetic variation, π and θ, were calculated using MEGA7 [73]. Assuming the sample from 123 infected *T. infestans* is representative of the *T. cruzi* population in vectors in Arequipa, the probability that a distinct lineage could be co-circulating but not detected by chance can be calculated using a binomial distribution. The minor lineage must constitute less than 2.41% of the total population in order for there to be a statistically significant chance that a distinct lineage was not detected in any of 123 sampled *T. cruzi.*

## Acknowledgements

The authors would like to thank Philippe Lemey for his advice regarding the use of Bayesian phylogenetics. The authors would also like to acknowledge Stephanie Seifert and Jill Devine for their assistance in the laboratory. The authors gratefully acknowledge the members of the Universidad Peruana Cayetano Heredia and the University of Pennsylvania Zoonotic Disease Research Lab in Arequipa, Peru, for their contributions, especially Carlos Condori and Luis Zamudio. The authors also thank Danitza Pamo, Jose Ylla, Jose Qusipe, Paul Picardo and Gabriela Bustamante for their contribution during the isolation and maintenance of the *T. cruzi* strains. In addition, the authors wish to acknowledge the support provided by the following institutions: Ministerio de Salud del Perú (MINSA), the Dirección General de Salud de las Personas (DGSP), the Estrategia Sanitaria Nacional de Prevención y Control de Enfermedades Metaxenicas y Otras Transmitidas por Vectores (ESNPCEMOTVS), the Dirección General de Salud Ambiental (DIGESA), the Gobierno Regional de Arequipa, the Gerencia Regional de Salud de Arequipa (GRSA), the PanAmerican Health Organization (PAHO/OPS) and the Canadian International Development Agency (CIDA).

---

**Figure S1.**
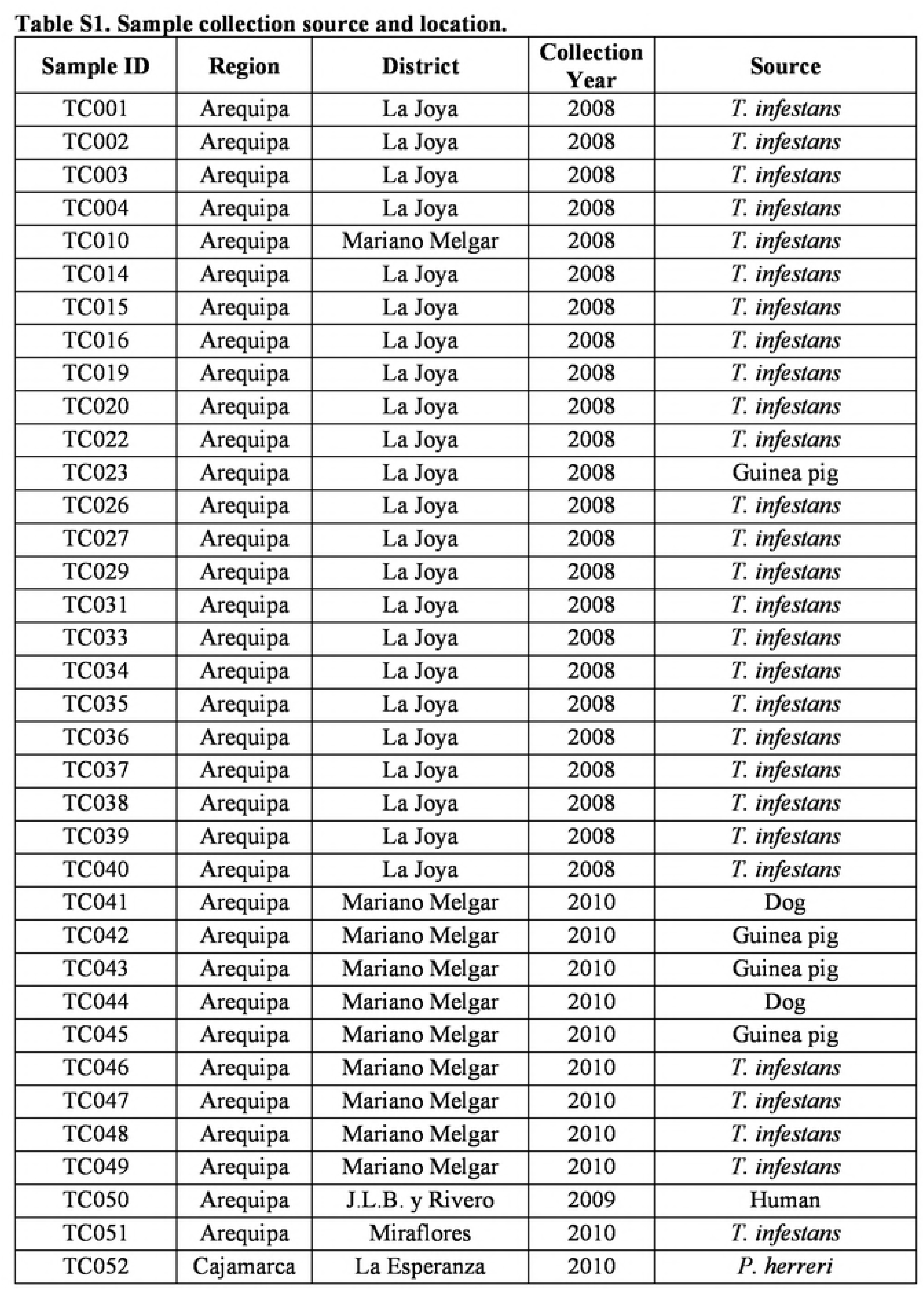
Sample collection locations and sources.

